# Comparative assessment of quantification methods for tumor tissue phosphoproteomics

**DOI:** 10.1101/2022.02.07.479414

**Authors:** Yang Zhang, Benjamin Dreyer, Natalia Govorukhina, Alexander M. Heberle, Saša Končarević, Christoph Krisp, Christiane A. Opitz, Pauline Pfänder, Rainer Bischoff, Hartmut Schlüter, Marcel Kwiatkowski, Kathrin Thedieck, Peter L. Horvatovich

## Abstract

With increasing sensitivity and accuracy in mass spectrometry, the tumor phosphoproteome is getting into reach. However, the selection of quantitation techniques best-suited to the biomedical question and diagnostic requirements remains a trial and error decision as no study has directly compared their performance for tumor tissue phosphoproteomics. We compared label-free quantification (LFQ), spike-in-SILAC (stable isotope labeling by amino acids in cell culture) and TMT isobaric tandem mass tags technology for quantitative phosphosite profiling in tumor tissue. TMT offered the lowest accuracy and the highest precision and robustness towards different phosphosite abundances and matrices. Spike-in-SILAC offered the best compromise between these features but suffered from a low phosphosite coverage. LFQ offered the lowest precision but the highest number of identifications. Both spike-in-SILAC and LFQ presented susceptibility to matrix effects. Match between run (MBR)-based analysis enhanced the phosphosite coverage across technical replicates in LFQ and spike-in-SILAC but further reduced the precision and robustness of quantification. The choice of quantitative methodology is critical for both study design such as sample size in sample groups and quantified phosphosites, and comparison of published cancer phosphoproteomes. Using ovarian cancer tissue as an example, our study builds a resource for the design and analysis of quantitative phosphoproteomic studies in cancer research and diagnostics.

## Introduction

Phosphorylation events mediated by oncogenic kinases are widely recognized as major drivers of tumorigenesis. Despite more than 50 kinase inhibitors approved for cancer treatment by Food and Drug Administration (FDA) and European Medicines Evaluation Agency (EMEA)^1^, immediate and acquired drug resistance remains a key medical challenge. Toward personalized treatments, genome and transcriptome analysis based detection of somatic mutations and mRNA changes has led to breakthroughs such as the identification of HER2-positivity that guides therapies with HER-targeting antibodies^2,3^ or BCR-ABL fusion indicating a response to Imatinib and its analogues^4^. Although these therapies significantly improve the survival rates e.g. in breast cancer and leukemia, they suffer from significant initial and acquired resistance^5舑7^. For most cancers and kinase-directed drug therapies reliable predictive markers of initial drug response and acquired resistance are largely missing. Pathway analysis in cancer cell lines highlights the potential to predict drug sensitivity^8^, and calls for translation to patient tumor tissues. Measuring the phosphorylation events and networks directly targeted by kinase inhibitors in the patient’s tumor may offer direct access to tumor- and patient-specific alterations governing drug resistance. Comprehensive quantitative coverage of the cancer- and patient-specific tumor phosphoproteome may identify patient subgroups with common response or escape mechanisms and may open new avenues to precision oncology by revealing a specific kinase signature in individual patients. Antibody-based techniques such as immunohistochemistry (IHC) and reverse-phase protein arrays (RPPA) are used in diagnostics and research, but they are limited by the availability of reliable antibodies and the low quantitative accuracy and robustness of these assays^9,10^. Mass spectrometry (MS)-based phosphoproteomic methods that combine the latest generation of instruments^11–14^ with advanced computational tools^15–18^ allow us to detect and quantify thousands of phosphosites. Label-free quantification (LFQ), stable isotope labeling by amino acids in cell culture (SILAC)^19^ and chemical labeling e.g. with isobaric tandem mass tags (TMT)^20^ or isobaric tags for relative and absolute quantitation (iTRAQ)^21^ are widely used for quantitative (phospho)proteomics and have been characterized in detail regarding their performance for studies in *in vitro* systems such as yeast or mammalian cell cultures^22–27^. But we do not yet know whether the results translate to clinical phosphosproteomics in tumors. Tumor tissue differs from cell cultures in many aspects that are relevant for sample preparation and MS analysis. For instance, tumor tissue is typically snap-frozen and must be powderized prior to lysis, whereas cultured cells can be directly taken up in lysis buffer. Tumor tissue also contains different cell types as well as extracellular matrix and is thus more heterogeneous than cultured cells^28^. In the last five years, a series of studies have used LFQ, spike-in-SILAC and TMT based quantification separately for tumor proteomics^29–34^ or phosphoproteomics^35–37^ highlighting the need for a comparative assessment of quantitative phosphoproteomic methodologies in tumor tissue. The classical SILAC approach requires full metabolic labeling of the entire proteome, and has been mainly limited to cell culture analysis with few exceptions^38^. Mann and colleagues introduced spike-in-SILAC^39^ whereby a SILAC-labelled reference cell line is used to generate thousands of isotopically labeled peptides as internal standards for tissue proteome quantification. Ovarian cancer is well-accessible to clinical proteomics and presents a high unmet medical need due to limited treatment options and short survival^40^. Using ovarian cancer as an example, we compared the performance of LFQ-, spike-in-SILAC - and TMT-based phosphoproteomics regarding accuracy, precision and robustness towards variation of the matrix and the phosphosite abundance. We use the term robustness in this article as the relative independence of phosphosites quantification from the number of replicates, phosphorylation levels and matrix effects. The SKOV3 cell line, derived from ascites of a human ovarian adenocarcinoma^41^ was used to compare our results from tumor tissue to a matched cell line.

### Experimental Section

#### SKOV3 cell culture and SILAC labeling

Experimental details of the cell culture conditions can be found in the ***Supporting Information***.

#### SKOV3 cell line and ovarian cancer tissue lysis

The detailed procedure of sample lysis can be found in the ***Supporting Information***.

#### Dephosphorylation and sample pooling

For dephosphorylation of the samples, it was required to exchange the protein lysis buffer with a phosphatase reaction buffer (10 mM Tris-HCl, 5 mM MgCl_2_, 100 mM KCl, 0.02% Triton X-100, pH 8.0), as alkaline phosphatases are inactive in the urea-containing protein lysis buffer. For this purpose, lysates of light labeled cells and tissues were loaded on a centrifugal filter unit (AMICON ULTRA-15 15ML - 10 KDa, cat. no. UFC901024) and centrifuged at 3,000 g, RT for 1 h. Subsequently, the centrifugal filter unit was washed five times with 5 ml phosphatase reaction buffer. To remove the phosphatase groups of the proteins^42,43^, alkaline phosphatase (protein: TSAP, 100:1, w/w, FastAP Thermosensitive Alkaline Phosphatase, cat. no. EF0651) was added to the lysates. Samples were incubated at 37°C for 1 h. After inactivation of the reaction at 74°C for 15 min, the buffer was changed back to the protein lysis buffer, as described before using the centrifugal filter unit. Subsequently, the protein concentrations were determined using the BCA Assay kit. The intensities of the phosphopeptides before and after phosphatase treatment were measured using LC-MS/MS. The dephosphorylation efficiency was calculated as the ratio of the total intensity of the phosphopeptides before and after phosphatase treatment. To achieve three sample groups with different phosphorylation levels, the samples with and without phosphatase treatment were pooled according to the scheme in **Figure S1**. The original non-treated lysates were mixed with the dephosphorylated samples lysate at ratios of 1:5 (1X), 1:1 (3X), and 1:0 (6X). Next, each sample was divided into six aliquots containing 1 mg for LFQ, 500 µg for spike-in-SILAC, and 300 µg for TMT, each. 500 µg of the heavy SILAC labelled cell lysate was mixed 1:1 with 500 µg of the 1X, 3X, or 6X tissue samples or the 1X, 3X, or 6X of the light SILAC-labelled cell line samples, respectively (Figure S1, panel 3).

#### Protein reduction, alkylation and digestion

Protein lysates were subjected to protein reduction, alkylation, and then trypsin digestion. The detailed experimental procedure can be found in the ***Supporting Information***.

#### TMT labeling

For TMT labeling the dried peptides were re-solubilized in 849 µL TEAB/ACN buffer (80 % H_2_O, 10% 1M TEAB, 10% ACN). The TMT (TMT, Thermo Scientific(tm), TMT10plex(tm) Isobaric Label Reagent Set, cat. no. 90406) reagents were solubilized in 100% ACN to a final concentration of 100 mM. 150 µL of the TMT stock reagents were added to the peptides according to **Figure S1** and incubated for 1 h at RT. To prevent side reactions, hydroxylamine (Thermo Scientific(tm), 50% Hydroxylamine for TMT experiments, cat. no. 90115) was added to a final concentration of 0.25% [w/v], followed by incubation for 15 min at RT. Next, samples were pooled according to the scheme in **Figure S1** and incubated for an additional 15 min. To reduce the concentration of ACN below 5% the samples were diluted 1:1 with 2% TFA, before dilution with H_2_O. Samples were desalted using SepPak tC18 cartridges (Sep-Pak tC18 1 cc Vac Cartridge, 200 mg Sorbent per Cartridge, 37 - 55 µm, cat. no. WAT054925). The desalting steps were performed as described before increasing the used buffer volumes to 2 mL. 50 µg of the eluent was used for the LC-MS/MS analysis and 2950 µg for the following IMAC enrichment. All the samples were dried using a SpeedVac.

#### Phosphopeptides enrichment

The phosphopeptides were enriched using the Immobilized metal affinity chromatography (IMAC) method (High-Select(tm) Fe-NTA Phosphopeptide Enrichment Kit, cat. no. A32992) according to the manufacturer protocol. In brief, the lyophilized peptide samples were suspended in 200 µL of IMAC Binding/Wash Buffer. After removing the bottoms closure of the IMAC spin columns and unscrewing the screw caps, the columns were centrifuged at 1000 × g for 30 sec to remove the storage buffer. For equilibration, the columns were washed twice with IMAC Binding/Wash Buffer. 200 µL of the suspended peptide samples were loaded on the equilibrated spin columns, and the columns were closed with the screw caps. The resin was mixed with the sample gently until the resin was in suspension. The suspension was incubated for 30 min, with gentle mixing every 10 min. Columns were washed three times with Binding/Wash Buffer and one time with HPLC grade H_2_O. The phosphopeptides were eluted by adding two times 100 µL of Elution Buffer to the column and centrifuged at 1000 × g for 30 sec. Samples were dried in a SpeedVac.

#### LC-MS/MS measurement and Raw data processing

For LC-MS/MS analysis, samples were injected on an ultrahigh performance nano liquid chromatography system (Dionex UltiMate 3000 RSLCnano, Thermo Scientific, Bremen, Germany) coupled to an Orbitrap mass spectrometer (Fusion(tm), Thermo Fisher Scientific) with a nano electrospray source. All LC-MS/MS data were processed with MaxQuant^15^ version 1.6.5. Detailed information of the LC-MS/MS separation and MS parameters can be found in the ***Supplemental Experimental Section***. All the Raw and MaxQuant processed data is available at ProteomeXchange under PXD030450 identifier. The identified phosphosite with quantitative values are available in Supporting information (**Table S1**).

#### Data pre-processing and statistics analysis

The pre-processed LC-MS/MS data were further processed and statistically analyzed using R 3.6.5^44^. The phosphosites intensity of LFQ and spike-in-SILAC was normalized to the total protein intensity^45^. The phosphosites intensity of TMT samples was normalized to the reference sample to remove the variance from batches^46^, before further normalization on the total protein intensity. For the correlation matrix heatmap, the correlation matrix of normalized intensity between each sample were firstly calculated and then plotted by ggplot2^47^. Linear regression model were applied to classify the sample groups and the performance of regression models are visualized by the ROC curve in ggplot2^47^. To visualize kinase substrate sites that were enriched among the quantified phosphosites, enrichment analysis of predictive kinases was performed. The kinase-substrate relationship was extracted from the thePhosphoSitePlus^48^ database. For the sequence motif analysis of the quantified phosphorylation sites, the ggseqlogo^49^ package was applied to generate the sequence logos. The probability of the amino acid around the identified phosphosites is shown in the sequence motif plot, with annotating the residue physicochemical properties by color. The barplot, distribution plot and violin plots were plotted using the ggplot2^47^ package. The VennDiagram^50^ package was used to plot Venn diagrams. R script used for data analysis and visualization is available on GitHub at https://github.com/functional-proteo-metabolomics/quantitative_phosphoproteomics.

## Results

### Sample preparation and MS measurements

The snap-frozen ovarian tumor tissue was powderized and taken up in ice-cold lysis buffer. SKOV3 cells were cultured in full medium. After a wash step, ice-cold lysis buffer was added directly to the tissue culture plates, and the cells were scraped off and transferred to tubes. Samples were sonicated and centrifuged, and the protein concentrations were adjusted to 1 mg/mL. All steps were performed on ice. To obtain samples with known ratios of phosphopeptide quantities (**Figure S1A, panel A**), we dephosphorylated tissue and cell lysates containing 20 mg protein with alkaline phosphatase. The dephosphorylation efficiency was assessed by LC-MS/MS. After alkaline phosphatase treatment, the overall intensity of identified phosphopeptides was reduced to 1.6% for SKOV3 cells and 3% for ovarian tumor tissue (**Figure S2A**). We mixed the non-treated lysate with the dephosphorylated lysate at ratios of 1:5 (termed in the following as sample 1X), 1:1 (3X), and 1:0 (6X) (**Figure S1B**). Thus, the sample with the lowest phosphosite quantity was 1X. The phosphosite quantity in the 3X sample was thrice as high, and it was 6 times as high in the 6X sample. This resulted in a known fold change of 2 (FC2) for the 6X versus the 3X sample. The 1X versus 3X samples exhibited a known FC of 3 (FC3), and the 1X versus 6X samples had a known FC of 6 (FC6). The known fold changes served as the ground truth based on which we assessed in the following the performance of spike-in-SILAC, TMT, and LFQ based phosphosite quantification. In order to assess the technical variability, we performed the experiments in six technical replicates, which were prepared separately from the lysate samples (**Figure S1**). For this purpose, each sample was divided into six aliquots of 1 mg for LFQ, 500 µg for SILAC, and 300 µg for TMT.

To generate the spike-in-SILAC standard, the SKOV3 cells were cultured for 5 passages in medium containing heavy lysine and arginine^51^. For the cell lysate to be compared with tissue sample, SKOV3 cells were cultured in light SILAC media. The labeling efficiency was >98% and the arginine-to-proline conversion was 1% or below (**Figure S2B, C**). 500 µg of the heavy SILAC labelled cell lysate was mixed 1:1 with 500 µg of the 1X, 3X, or 6X tissue lysates or of the 1X, 3X, or 6X light SILAC-labelled cell lysates, respectively (**Figure S1 B, C**), resulting in a total amount of 1 mg for each sample. All samples were reduced, alkylated, digested with trypsin for 16 h, and desalted using reversed phase solid phase extraction. TMT 10-plex was used to label 6 technical replicates of the 6X, 3X, and 1X tissue and cell line samples, which we assigned to two batches (**Figure S1C**). Each batch contained in addition a reference sample that was composed of 100 µg of the 6X, the 3X, and the 1X sample. For TMT analysis, two TMT analysis batches were prepared, each consisting of 10 TMT-channels composed of 9 samples and one reference sample **Figure S1C (bottom)**. For each channel, 300 µg of TMT-derivatized sample was used. This results in a total sample amount of 3 mg per TMT batch. Thus, we stayed below the maximum binding capacity of the Fe-NTA (nitrilotriacetic acid)-based IMAC columns (5 mg). Before IMAC enrichment, 50 µg of each sample were set apart for total protein concentration analysis. The rest of the samples were subjected to phosphopeptide enrichment by IMAC. All samples were dried and analyzed by LC-MS/MS using an Orbitrap Fusion instrument. The gradient length was kept the same for all samples (2 h). The LFQ and SILAC samples were analyzed by data dependent acquisition (DDA)^52^ tandem MS (MS2). The TMT samples were analyzed by synchronous precursor selection MS3 (SPS-MS3)^53,54^, to minimize reporter ion cross-contamination from peptides co-isolated for fragmentation in the same isolation window. The data was analysed using the MaxQuant^15^ suite. Shotgun proteomics by DDA is a widely used technology^52^. Although DDA achieves fast and accurate large-scale proteome analysis, missing values of 50% or higher are inherent characteristics of the stochastic precursor selection^55,56^. Data pre-processing with match between runs (MBR) is a common approach to reduce the number of missing values in DDA and perform identification transfer by matching non-identified single-stage MS (MS1) features to identified ones^27,57^. We compared data processing without and with the MBR^58,59^ option for LFQ and spike-in-SILAC, including the requantify (req) option that is typically applied in SILAC experiments to extract the signal of non-detected peaks, which are complementary to successfully identified light or heavy SILAC singleton peaks^15^.

### Phosphosite and phosphopeptide identifications

We compared the total measurement time, the number of recorded MS2 spectra and MS3 spectra, the number of identified MS2 spectra and the number of identified phosphosites between LFQ, SILAC and TMT (**Table S1**). In TMT, the MS3 spectra were used for quantification and the MS2 spectra were used for identification. Therefore, we compared the number of acquired and identified MS2 spectra to assess the identification performance. The threshold for the phosphosite localization probability was set to 0.75^58^.

For tumor tissue, SILAC provided the highest number of detected (n_spectra_=613,676) and identified (n_id_=240,688) MS2 spectra, followed by LFQ (n_spectra_=471,216, n_id_=171,752) and TMT (n_spectra_= 63,421 and n_id_=8,122). However, the number of phosphopeptide identifications was higher with LFQ (8,932) than with SILAC (6,923). The discrepancy between the number of MS2 spectra and the number of identified phosphopeptides may be explained by the SILAC pair-induced increased complexity at the MS1 level. The isolation and fragmentation of the light and heavy phosphopeptide leads to a redundant identification, which reduces the number of identified phosphopeptides compared to LFQ^60^. TMT resulted in the lowest number of identified phosphopeptides (n= 4,281). Similar observations were made for cell lysate (**Table S1**).

In line with the number of identified phosphopeptides, LFQ provided the highest number of identified phosphosites (n=5,812) followed by SILAC (n=4,828) and TMT (n=3,578) for tumor tissue. Similar results were obtained for cell lysate (**Table S2**). 1,640 (20%) and 1,205 (18.6%) of the phosphosites were identified by all three methods in the tumor and cell line samples, respectively (**Figure 1A and B**). SILAC and LFQ showed an overlap of ca. 40% (40.2% for tumor tissue, 43.2% for cell line). Lower overlap was observed between TMT and LFQ or SILAC for both tissue (TMT-LFQ: 25.9%, TMT-SILAC: 21%) and cell line (TMT-LFQ: 20.7%, TMT-SILAC: 21.7%). This may be explained by TMT-derivatization due to which the resulting phosphopeptides differ in their chemical composition and physico-chemical properties^20^ from phosphopeptides analyzed by LFQ and SILAC that are chemically equivalent and differ only in their isotopic composition. We compared the frequency of b- and y-fragment ions of all identified phosphosites (**Figure 1C, D**). LFQ and SILAC exhibited similar fragment ion frequencies in both tissue and cell line samples, with a higher frequency of y-ions (up to 75%) than b-ions (up to 25%), as expected for non-derivatized tryptic peptides^61^. For TMT-derivatized phosphopeptides, the frequency of y- and b-fragment ions was equal. Similar results were obtained for the non-phosphorylated peptides in the input sample (**Figure S3A, B**) which is in agreement with earlier observations by Shen et al^62^. The higher frequency of b-ions may be explained by the introduction of a tertiary amine at the N-terminus as a result of TMT labeling. The tertiary amine of the TMT-tag has a higher gas phase basicity than the primary amine of the N-terminus of a tryptic peptide. As a result, the proton at the piperidine ring of the TMT-tag linked to the peptide N-terminus is less mobile, and b- and y-fragment ions at are generated at similar frequencies^63^. The TMT-label impacts both the ionization and the fragmentation of the derivatized phosphopeptide^64^, which may explain the low overlap of identifications in TMT with LFQ and spike-in-SILAC. Our results illustrate that a derivatization strategy can shift the identifications to different subpopulations of phosphosites and peptides, which needs to be considered when comparing datasets acquired without and with chemical labelling.

**Figure 1:**
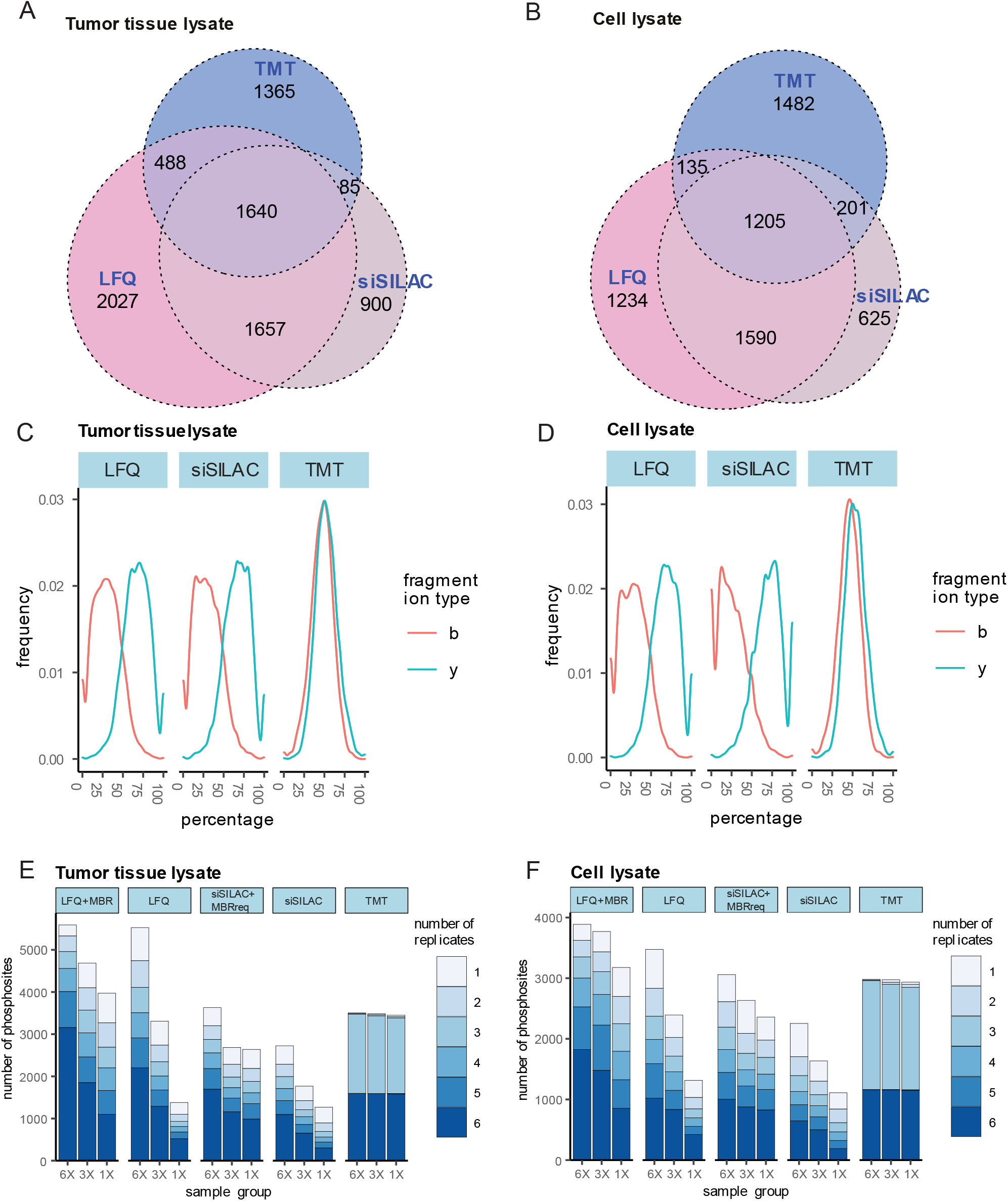
Comparison of phosphosite identifications by LFQ, spike-in-SILAC and TMT in tumor tissue lysate or cell lysate samples. A, B: Venn diagram showing the number of identified phosphosites for the three different quantification methods (LFQ, spike-in-SILAC and TMT) in the tumor tissue lysates (A) and cell lysates (B). C, D: Density plots showing for each quantification method the distribution of b- and y-ions of the identified phosphopeptides in tumor tissue lysates (C) and cell lysates (D). E, F: Bar plots showing the number of phosphosites identified in each sample group (6X, 3X and 1X) for the different quantification methods used in tumor tissue lysates (E) and cell lysates (F). The color intensity indicates the number of replicates in which the phosphosites were identified. Only phosphosites with a localization probability of at least 0.75 were considered for the analysis.

We next compared the reproducibility of phosphosite identifications across the six technical replicates in tumor tissue (**Figure 1E**). The LFQ and SILAC data sets were processed with or without activation of the MBR feature in MaxQuant. The reproducibility of the phosphosite identification is reflected by the distribution of the numbers of identified phosphosites across the replicates. MBR can be considered as identification transfer step and therefore influences the reproducibly of phosphosite identification. For TMT, MBR has recently been implemented for MS2 methods^65^. To enhance the quantification accuracy, we opted for SPS (synchronous precursor selection)–MS3 technology^53,54^. As expected, MBR increased the number of reproducibly identified phosphosites for both LFQ and SILAC. The number of phosphosites identified across six replicates decreased with decreasing phosphosite quantity (i.e., 6X vs. 3X vs. 1X) for LFQ and spike-in-SILAC with and without MBR. This indicates that with LFQ and spike-in-SILAC reproducible identification increases with phosphosite quantity and the resulting signal intensity. This was in contrast to TMT for which the number of phosphosites reproducibly identified across 6 replicates was constant over the entire range of phosphorylation quantity (n=1593 in 6X, n=1591 in 3X and n=1582 in 1X). This likely comes from the multiplexing of the 1X, 3X and 6X samples due to which low intensity signals in the 1X sample can be identified based on the higher average intensity from the pooled 1X, 3X and 6X samples^20^. The higher average phosphosite quantity and signal intensity in the pooled TMT sample also explains the better reproducibility of identifications for samples with low phosphosite quantity (i.e. 1X and 3X) as compared to LFQ and spike-in-SILAC. For the high phosphosite quantity (6X) TMT yielded also more reproducibly identified phosphosites than spike-in-SILAC, but the MBR option increased this number beyond that of TMT. LFQ without and with MBR yielded higher numbers of reproducible identifications than TMT for high phosphosite quantities (6X). The reproducible identification of phosphosites in six or three replicates in TMT (with other replicate numbers largely missing) arises from the fact that the TMT samples were measured in two batches with three replicates per batch (**Figure S1C**). Therefore, a given phosphosite was identified in either of the batches (i.e. in 3 replicates), or in both batches (i.e. in 6 replicates). Comparable results were obtained for the reproducibility of the identification of phosphopeptides, with higher overall numbers as phosphosites can be assigned to several peptides (**Figure S4A**).

The overall numbers of reproducibly identified phosphosites and phosphopeptides from cell lysate were on average lower than from tissue lysate (**Figures 1F, S4B**). Also here, TMT yielded a similar number of reproducible identifications across the different phosphorylation quantities. TMT yielded a higher number of reproducible identifications than spike-in-SILAC without and with MBR and LFQ across all phosphosite quantities. Only LFQ with MBR in a sample with a high phosphosite quantity (6X) performed better than TMT in terms of reproducibly identified phosphorylations. In both matrices, tumor tissue und cultured cells, TMT exhibited better robustness than LFQ and spike-in-SILAC regarding reproducibility of identifications across the whole range of phosphosite quantities. In tumor tissue with high phosphorylation quantity, spike-in-SILAC and LFQ outperformed TMT regarding the number of reproducible identifications, but this advantage was lost for lower phosphosite quantities.

### Reproducibility, accuracy and precision of phosphosite quantification

We assessed the reproducibility of phosphosite quantification by calculating the coefficient of variation (CV) distribution (**Figures 2A, B and S5A-D**). We observed no major difference in CV distribution in datasets analyzed with or without MBR both for LFQ (**Figure S5A, B**) or spike-in-SILAC (**Figure S5C, D**). TMT provided the lowest CVs, followed by spike-in-SILAC+MBRreq and LFQ+MBR (**Figure 2A, B)**. For TMT the CV distribution was similar across phosphosite quantities (1X, 3X, 6X). Thus, TMT exhibits the highest reproducibility for quantification of low-abundant phosphosites. TMT multiplexing reduces the technical variability as multiple samples are pooled and processed together during sample preparation (e.g. phosphopeptide enrichment and desalting by solid-phase extraction) and LC-MS/MS measurements. A reference channel in each TMT-batch further reduced the measurement variability between the two batches. In spike-in-SILAC, the spiked-in heavy-labeled phosphopeptides serve a similar purpose as the TMT reference channel, namely to reduce variability introduced during sample preparation and LC-MS/MS analysis. This likely explains the lower CV values with spike-in-SILAC as compared to LFQ, for which all experimental and measurement steps were conducted independently and no reference sample was spiked-in. TMT shows lower CVs compared to spike-in-SILAC, although mixing of the multiplexed samples in spike-in-SILAC occurs earlier than in TMT. The smaller CVs of TMT are most likely due to the peak quantification in smoothed MS3-orbitrap mass spectra, whereas spike-in-SILAC used MS1-based 2D peak quantification in an LC-MS map. In these, the chromatographic peaks have a higher noise level than the signals in smoothed MS3-orbitrap mass spectra^66^. This may enable a more precise quantification in TMT data as compared to spike-in-SILAC data.

**Figure 2:**
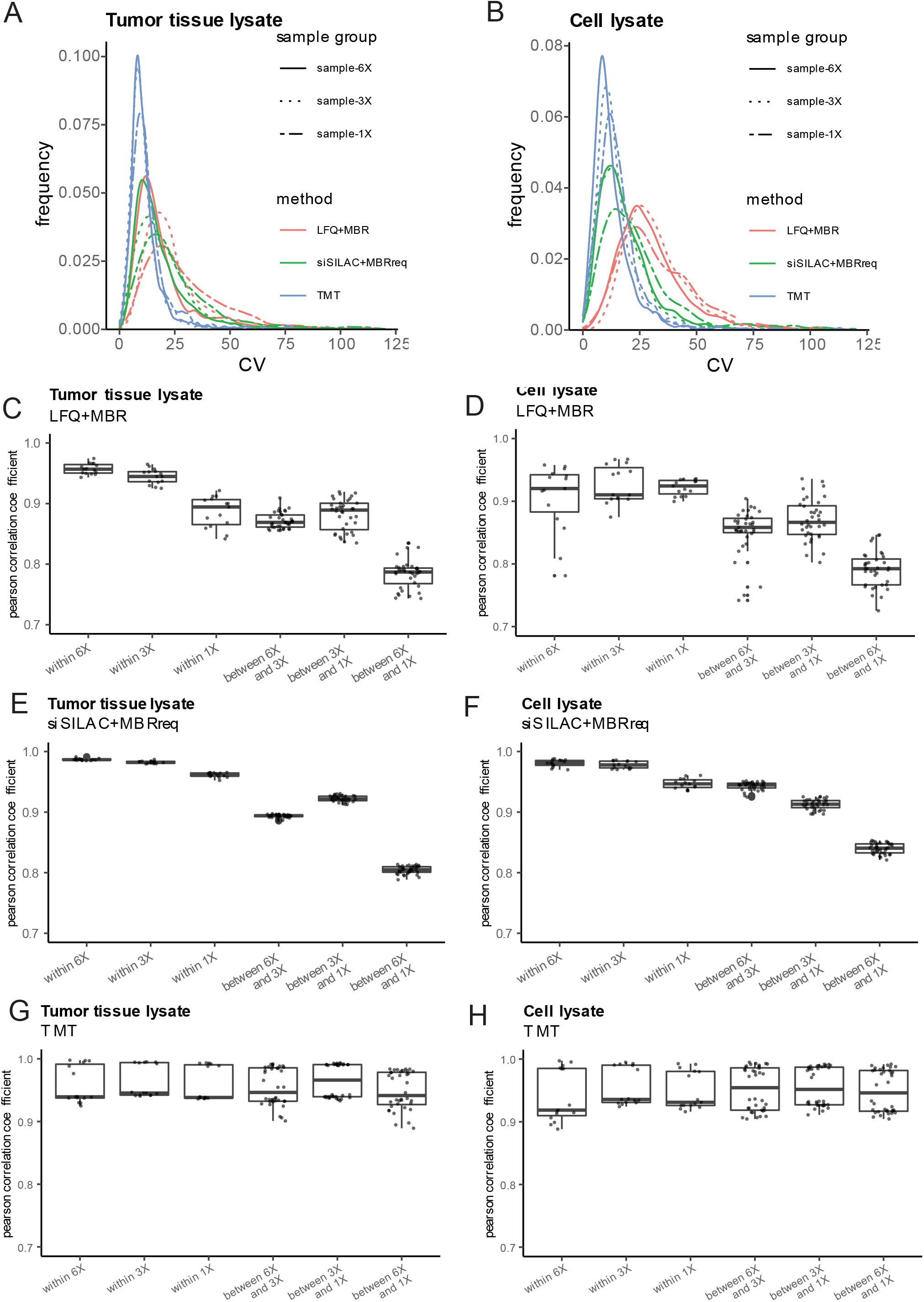
Evaluation of the reproducibility of phosphosite quantification. A, B: Density plots showing the distribution of CV values for phosphosites quantified with LFQ+MBR, spike-in-SILAC+MBRreq and TMT in the tumor tissue lysates (A) and cell lysates (B). Different line types (solid, dotted, dashed) indicate different sample groups (6X, 3X and 1X). C-H: Boxplots visualizing the pairwise correlation coefficients within the same sample group (i.e. 6X vs. 6X, 3X vs. 3X, 1X vs. 1X) and between the different sample groups (i.e., 6X vs. 3X, 3X vs. 1X, 6X vs. 1X) for each method in tumor tissue lysates (C: LFQ+MBR, E: spike-in-SILAC+MBRreq, G: TMT) and cell lysates (D: LFQ+MBR, F: spike-in-SILAC+MBRreq, H: TMT). The boxes show the first, second (median) and third quartile.

The CV distribution reflects the variability based on the mean of six replicates for each phosphosite. To assess the agreement between datasets in a pairwise manner, we analyzed the linear correlation between two datasets by calculating the pairwise correlation of phosphosite quantities for phosphopeptides identified and quantified in both samples (**Figures 2C-H and S5E-H**). Similar to the CV plots, no major difference was observed in the correlation heatmaps with and without MBR/MBRreq for LFQ and spike-in-SILAC (**Figures 2C-F and S5E-H**). For tissue and cell lysates, both TMT (**Figure2G, H**) and spike-in-SILAC (**Figure 2E, F**) showed Pearson correlation coefficients above 0.9 within a sample group for all phosphosite quantities (1X, 3X, 6X). In TMT, the correlation was always above 0.9 and did not change between different phosphosite quantities, although a batch effect was visible that broadened the correlation coefficient distribution. For spike-in-SILAC, lower coefficients were observed for low phosphosite quantities (1X). This was even more pronounced for LFQ, resulting in a Pearson correlation coefficient below 0.9 for tumor lysate with low abundant phosphosites (1X, **Figure 2C**). For all comparisons between sample groups (i.e., 6X vs. 3X, 3X vs. 1X, 6X vs. 1X), TMT yielded Pearson correlation coefficients above 0.9 (**Figure 2G, H**), reflecting the high reproducibility of TMT across different phosphosite quantities. For comparisons between sample groups, spike-in-SILAC and LFQ showed considerably lower Pearson correlation coefficients than TMT. The correlation coefficients were particularly low when samples with low phosphosite quantity (1X) were part of the comparison, reflecting the drop in reproducibility and higher error in the quantification of low abundant peaks for spike-in-SILAC and LFQ as the phosphosite quantity decreased. In summary, TMT exhibited the lowest variability and highest correlation for tissue and cell lysates, especially when the phosphosites were low-abundant.

### Accuracy and precision of the quantification of phosphosite fold changes

We evaluated the precision and accuracy of the determination of the fold-changes between samples with known relative phosphosite quantities (1X, 3X, 6X) (**Figure 3**). We assessed the distribution of detected fold changes relative to the theoretical fold changes (FC2, FC3 and FC6) by violin plots (**Figure 3A, B**) and by mean squared errors (MSE)^67^ of the sum of positive deviation or variance (**Figure 3C, D**), indicative of the quantification error in accuracy and precision, respectively. LFQ and spike-in-SILAC determined all fold-changes with high accuracy and yielded more accurate quantifications than TMT in tumor and tissue lysates (**Figure 3A-D**). The use of MBR decreased the accuracy (**Figure 3C, D**). MBR also reduced the precision of all determined fold changes, and this effect was more pronounced in spike-in-SILAC than in LFQ. Matched features with low abundance or false feature matching may be reasons for lower precision and accuracy in MBR^66^. This shows that MBR trades off a smaller number of missing values for accuracy and precision. TMT showed the highest precision and the lowest accuracy, consistently underestimating the expected fold change (**Figure 3A-D**).

**Figure 3:**
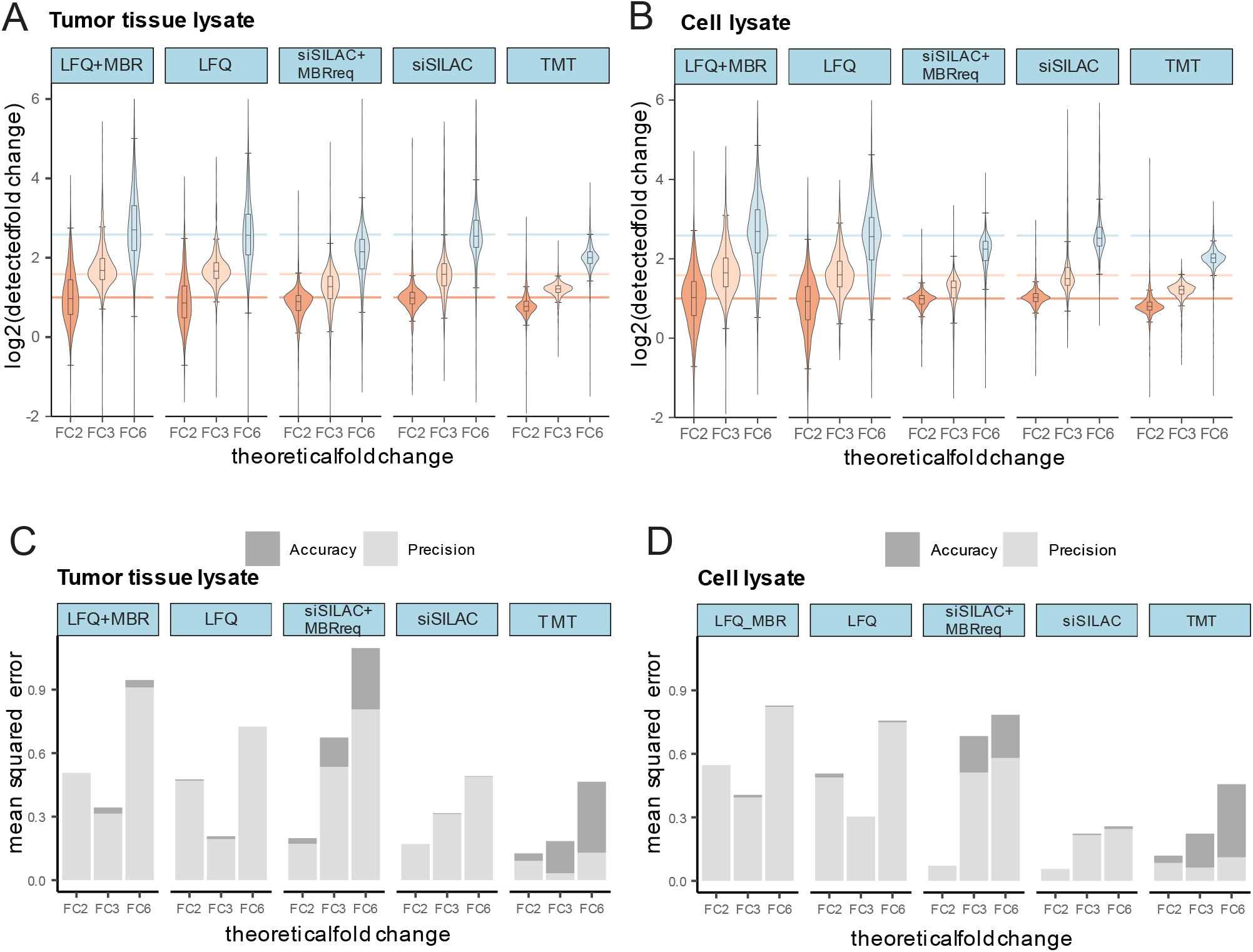
Evaluation of precision and accuracy of phosphosite quantification. (A, C) tumor tissue lysates. (B, D) cell lysates. A, B: Violin plots showing log_2_-transformed fold changes to evaluate quantification precision and accuracy errors of the methods. The boxes show the first, second (median) and third quartile. Whiskers show the minimum/maximum value within the 1.5 interquartile range. Expected log_2_-transformed fold changes were highlighted by colored lines. C, D: Bar plots showing mean squared errors of accuracy and precision. Mean squared errors were calculated as the sum of the square of positive deviation and variance for each method and all replicates.

We also investigated the performance in detecting significant differences in the abundance of phosphosites between the different fold changes. Phosphosites had to be quantified in at least three out of six replicates. Statistical analysis was performed using a two-sided t-test and Benjamini-Hochberg procedure to correct for multiple testing. Across all fold-changes, TMT determined the highest number of phosphosites as being significantly different for both tissue (**Figure 4A**) and cell (**Figure 4B**) lysates. The next-best results were yielded by LFQ+MBR and spike-in-SILAC+MBR. LFQ and spike-in-SILAC performed worst. As multiplexing in TMT reduces the issue of missing values, almost all identified phosphosites (> 98% for all sample groups) could be used for differential analysis. As stated earlier, also quantification based on smoothed MS3-orbitrap mass spectra yields more precise quantification and lower CVs, likely enhancing the discrimination of small differences. Spike-in-SILAC and LFQ resulted in a considerable proportion of phosphosites identified in less than three replicates, which prevented them from being used for differential analysis. The use of MBR in LFQ and spike-in-SILAC increased the number of significantly different phosphosites, in particular for the higher fold changes (FC3, FC6), which could be detected despite the lower precision (**Figure 3C, D**).

**Figure 4:**
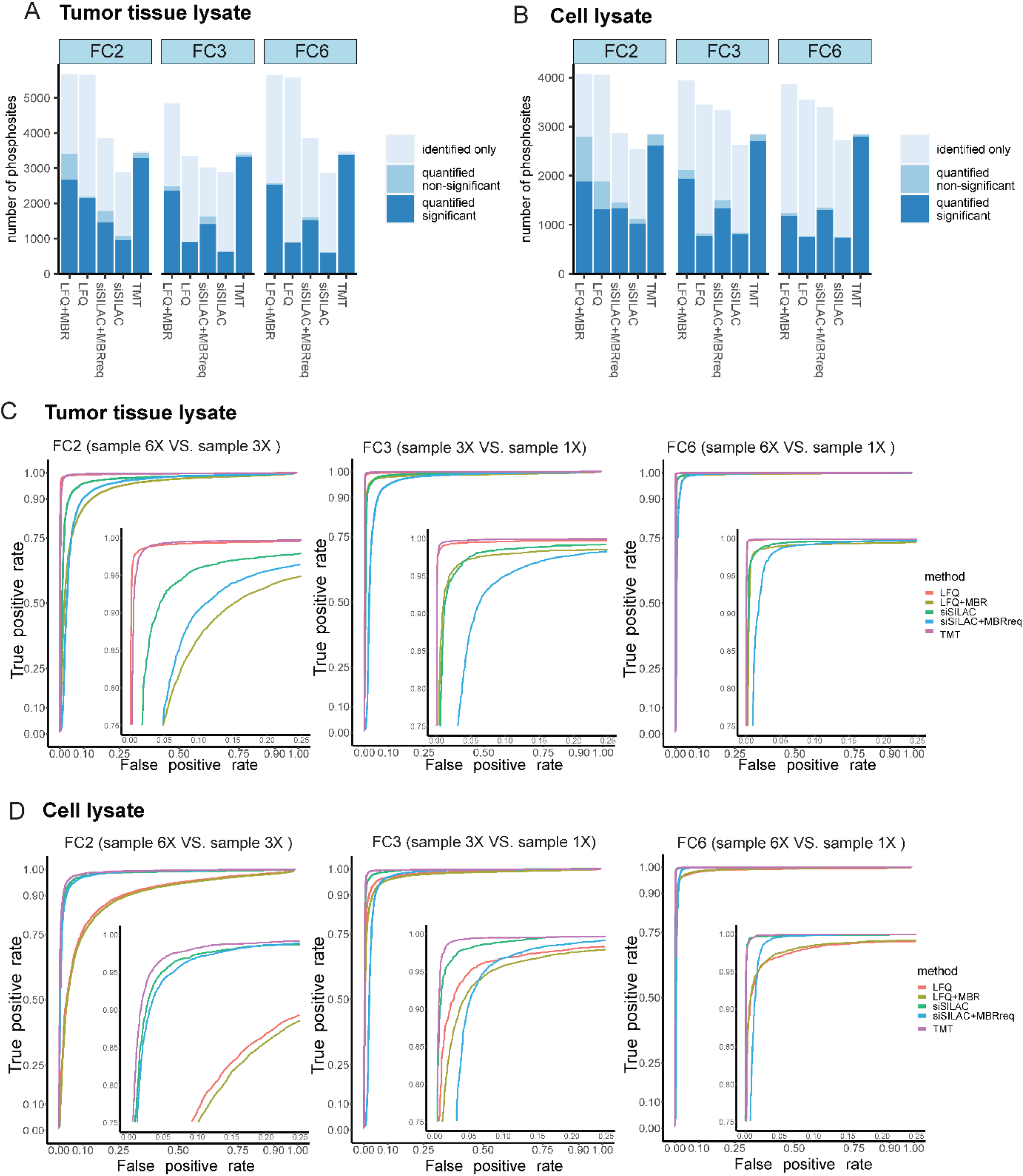
Performance in detecting quantitative phosphosite differences. A, B: Barplots showing the number of identified and quantified phosphosites in tumor tissue lysates (A) and cell lysates (B). Identified only: phosphosites identified in less than three replicates. Quantified non-significant: phosphosites identified and quantified in at least three out of six replicates, with FDR > 0.05. Quantified significant: phosphosites identified and quantified in at least three out of six replicates, FDR ≤ 0.05. C, D: Receiver operating characteristic (ROC) curves for evaluating the ability to diagnose the different phosphorylation quantities in tumor tissue lysates (C) and cell lysates (D). Zoomed-in ROC curves are presented in the lower right corner of each graph.

We calculated true-positive-rates (TPR) and false-positive-rates (FPR) of the differentially quantified phosphosites for each quantification method and visualized them in a receiver operating characteristic (ROC) curve^68^ (**Figure 4C, D**). We also calculated the area under the ROC (AUROC), which provide classification performance of TPR and FPR for whole thresholds range (**Table S3**). At an FPR threshold of 0.05, the TPR was above 0.95 for all quantification approaches and the largest fold-change (FC6), indicating that all methods were able to correctly determine a fold-change of six (FC6) between the phosphorylation quantities (**Fig. 4C, Table S3**). These results were also reflected by the AUROC values showing value higher than 0.98 for all methods for both tissue and SKOV3 cell lines (**Table S3**). In general, TMT showed the highest TPRs and AUROCs for all fold-changes (**Table S3**). For the tissue lysate and the smallest fold-change (FC2), TMT (TPR: 0.99) and LFQ (TPR: 0.99) showed a TPR higher than 0.95. In contrast, TMT (TPR: 0.97) and spike-in-SILAC (TPR: 0.95) showed the highest TPRs for cell lysate, whereas LFQ yielded TPRs below 0.95 for FC2 (TPR: 0.59) and FC3 (TPR: 0.92). AUROC values showed similar trends then TPR at FPR level of 0.05 (**Table S3**). MBR or MBRreq generally resulted in lower TPRs and AUROCs, further highlighting that MBR/MBRreq traded the numbers of identified phosphosites (**Figure 1E, F**) and of significantly different phosphosites (**Figure 4A, B**), for accuracy, precision (**Figure 3C, D**), and robustness of the quantification (**Figure 4C, D**).

Taken together, TMT showed the lowest accuracy but the highest precision and the highest number of significantly differential phosphosites for all fold-changes in tumor and cell lysate. TMT also provided the highest TPR regardless of sample matrix. In contrast, the matrix affected the TPRs for LFQ and spike-in-SILAC, with spike-in-SILAC exhibiting a TPR below 0.95 for tissue and LFQ yielding a TPR below 0.95 for cell lysate.

### Coverage of kinase targets by the quantified phosphosites

As TMT differs from LFQ and spike-in-SILAC regarding the identified phosphosite profiles (**Figure 1A, B**), we asked whether different quantification methods also introduce a bias in the coverage of quantified kinase substrate sites. For this purpose, an enrichment analysis^69^ for kinase substrates assigned to major oncogenic signaling pathways was conducted using the kinase-substrates database from PhosphoSitePlus^48^ (**Figure S6A, B**). TMT showed a higher coverage of most pathways, as compared to LFQ and spike-in-SILAC. This was mitigated by MBR and MBRreq, likely due to the higher number of quantified phosphosites through identification transfer from samples with higher phosphorylation quantities (**Figure 4A, B**).

However, only 10% of the quantified phosphosites were annotated in the kinase substrate database, limiting the power of this analysis and raising the possibility that further biases may have been missed. We therefore analyzed the coverage of kinase target motifs among the significantly quantified phosphosite. Whereas LFQ and spike-in-SILAC without or with MBR/MBRreq covered a similar array of kinase target motifs, their coverage differed for TMT (**Figure S7A, B**). We conclude that the profiles of both identified and quantified phosphosites differ between LFQ, spike-in-SILAC and TMT.

## Discussion

We report the first comprehensive comparative analysis of the quantitative performance of LFQ, spike-in-SILAC and TMT for the analysis of the tumor tissue phosphoproteome.

In summary, LFQ yielded the highest number of phosphosite identifications. MBR and MBRreq increased the number of identified phosphosites for LFQ and spike-in-SILAC (**Figure 1**). The lower number of phosphosites identified in TMT is likely explained by the SPS-MS3 approach where MS2 was used for phosphopeptide identification and phosphosite localization, while MS3 for quantification of the reporter ions requires a longer duty cycle and ion trap resonance CID. In ion-trap resonance CID used for TMT analysis, the depth of the potential prevents ion ejection and precursors can only be excited to a few electron volts, requiring long activation times to build up sufficiently high internal energy for fragmentation^70^. Gas-phase rearrangement reactions can occur prior to dissociation and have been reported for phosphate moieties of phosphoserine- and phosphothreonine-containing peptides^71^. Furthermore, only the precursor is excited in ion-resonance CID^72^. For phosphopeptides containing phosphoserines and phosphothreonines, this can result in abundant non-sequence informative fragment ions corresponding to the neutral loss of the phosphate moiety from the precursor^73,74^. In contrast, in beam-type HCD, which was used for fragmentation in LFQ and spike-in-SILAC, all ions are activated, and fragments including the phosphate neutral loss can undergo several consecutive fragmentation events^75^. Therefore, Beam-type HCD spectra contain more sequence informative fragment ions than resonance CID spectra, and the shorter activation times reduced potential gas-phase rearrangements. As beam-type HCD spectra are better suited for accurate phosphosite identification than resonance CID^76^, TMT yields in total less identifications than LFQ or spike-in-SILAC.

Nevertheless, TMT provided the highest number of reproducible identifications across all fold-changes and phosphosite quantities, whereas for LFQ and spike-in-SILAC the number of reproducibly identified phosphosites decreased with decreasing phosphosite quantities, which is mainly related to DDA precursor selection stochasticity. This suggests that in these approaches, reproducible identification is highly dependent on phosphosite quantities. The isobaric and multiplexing nature of TMT tags increases the signal intensity of phosphorylated peptides at the MS1 level, reducing the number of missing values. Although the number of reproducibly identified phosphosites could be increased with MBR and MBRreq for LFQ and spike-in-SILAC, TMT outperformed the other approaches in this respect for the analysis of both tissue and cell lysates.

Regarding phosphosite quantification, LFQ without MBR provided the highest accuracy and the lowest precision, whereas TMT showed the lowest accuracy and the highest precision, irrespective of the matrix (**Figure 3)**. MBR and MBRreq sacrifice precision and accuracy for a higher number of quantified phosphosites, with spike-in-SILAC suffering more severely from this issue than LFQ. In both tissue and cell lysate TMT exhibited the highest TPRs for the differentially quantified phosphosites (**Figure 4, Table S3**). In contrast, we found matrix effects for both LFQ and spike-in-SILAC. LFQ yielded high TPRs for the phosphoproteome in tissue but not cell lysate, whereas spike-in-SILAC showed high TPRs in cell lysate but not for tumor tissue. MBR decreased the TPRs and could not mitigate these matrix-effects. However, MBR increased the number of significantly different quantified phosphosites in LFQ and spike-in-SILAC, albeit not to the same amount as reached by TMT. This further highlights that MBR trades the number of phosphosites identified and quantified for quantification accuracy, precision, and robustness.

We analyzed the tumor phosphoproteome side-by-side with a matched cancer cell line to ensure comparability of our data with previous studies on cell lines and assess the robustness of the quantitative methods towards matrices as different as tumor and cell lysate. Our analysis of the cell lysate reflects overall the findings of Hogrebe et al. ^26^, although there are some differences in study design that need to be considered. Hogrebe et al. ^26^ used the protease Lys-C whereas we used trypsin as the most widely used protease for bottom-up proteomics^77,78^. TMT labeling after Titanium dioxide (TiO_2_) enrichment, as done by Hogrebe et al. ^26^, may reduce the advantage of TMT as multiplexing occurs late in the protocol and does not compensate technical variability introduced in the enrichment step. Therefore, we performed TMT labeling and multiplexing directly after trypsin digestion and used IMAC instead of TiO_2_ due to its higher selectivity, identification numbers and quantitative reproducibility^79^.

The quantification method significantly determines the study result in terms of identified, quantified and differentially assigned phosphosites, as well as quantification accuracy and last but not least the tumor related kinase phosphosite target profiles covered. For study design, the choice of the quantification method depends on the study aims. Although TMT outperformed spike-in-SILAC regarding quantification precision and was least amenable to matrix effects, it exhibited the lowest accuracy. If accuracy is a key requirement, TMT is therefore not the method of choice. Spike-in-SILAC provided the best compromise between accuracy, precision and robustness towards differences in phosphosite quantities. However, spike-in-SILAC suffered from a low phosphosite coverage and did not reliably detect low fold changes between phosphosite quantities in tissue. Its performance should therefore be assessed for the matrix to be analyzed (i.e. the tumor tissue) in order to determine the suitability of spike-in-SILAC regarding the TPR of low fold changes. LFQ was susceptible to matrix effects but exhibited a higher TPR in tumor tissue than in cell lysate. However, LFQ offered low precision, which was not counterbalanced by MBR. Rather, MBR degraded the precision and robustness of quantification. Recently, Bekker-Jensen et al. showed that data-independent acquisition (DIA) LFQ may overcome some of the limitations in DDA LFQ, which we observed in our study^80^. Also, practical considerations such as instrument time may influence the choice of method. TMT allows multiplexing which reduced the measuring time in our study by a factor of nine as compared to the LFQ and spike-in-SILAC which represents an inherent advantage of TMT regarding time and costs.

In conclusion, different quantification methods offer the highest accuracy, precision and phosphosite coverage in tumor tissue proteomics and thus the choice of quantification method is critical. The different behavior of TMT, LFQ and spike-in-SILAC as well as the influence of MBR should be also taken into account when comparing published tumor tissue phosphoproteomes. We advocate careful annotation of tissue phosphoproteomes to enable meaningful comparison.

## Supporting information

Supporting information (Supplementary figures, tables and experimental sections)

## Supporting information

The Supplemental_figures_tables_experimental_section.pdf contains supplementary **Figures S1-7, Tables S2-3** and supplementary experimental section. The “Table S1_Quantitative_Phosphosites table.xlsx” file contains the list of identified phosphosites with quantitative information (**Table S1**).

## Acknowledgments

We thank Ulrike Rehbein and Alienke van Pijkeren for critical reading of the manuscript.

We acknowledge support from the PROMETOV project that is supported by the national funding organizations and the EC under the framework of the ERA-NET TRANSCAN-2 initiative (to C.O., P.H., K.T); the MESI-STRAT project (grant agreement 754688 to K.T. and C.O.) and oLiMeR Innovative Training Network (Marie Skłodowska-Curie grant agreement 812616 to K.T.) which both received funding from the European Union Horizon 2020 Research and Innovation Program; the German Tuberous Sclerosis Foundation (to K.T.); Stichting TSC Fonds (to K.T.); the German Research Foundation (TH 1358/3-1 to K.T.); a Rosalind Franklin Fellowship of the University of Groningen (to K.T.); the Netherlands X-omics Initiative (NWO, project 184.034.019 to P.H.), the European Respiratory Society (ERS, RESPIRE3, project reference: R3201703-00121, to M.K.), the University of Innsbruck (project no: 316826, to M.K.) the Tyrolian Research Fund (project no: 18903, to M.K.) and the Deutsche Forschungsgemeinschaft (DFG) (INST 337/15-1, INST 337/16-1 &INST 152/837-1, to H.S.). Apart from the first author and the corresponding authors, all other authors are listed in alphabetical order.

## Author contributions

Study design and concept was made by Y.Z., N.G., M.K., K.T., P.H., sample preparation was performed by Y.Z. and LC-MS/MS analysis by Y.Z., B.D., C.K., the bioinformatics analysis was performed by Y.Z., P.H., clinical tissue collection was performed by C.A.O., P.P., project supervision and progress discussion were made by A.M.H., N.G., S.K., H.S., C.A.O., R.B., M.K., K.T., P.H., funding acquisition was made by S.K., H.S., C.A.O., K.T., P.H. The paper was written and revised by all authors.

## For Table of Contents Only

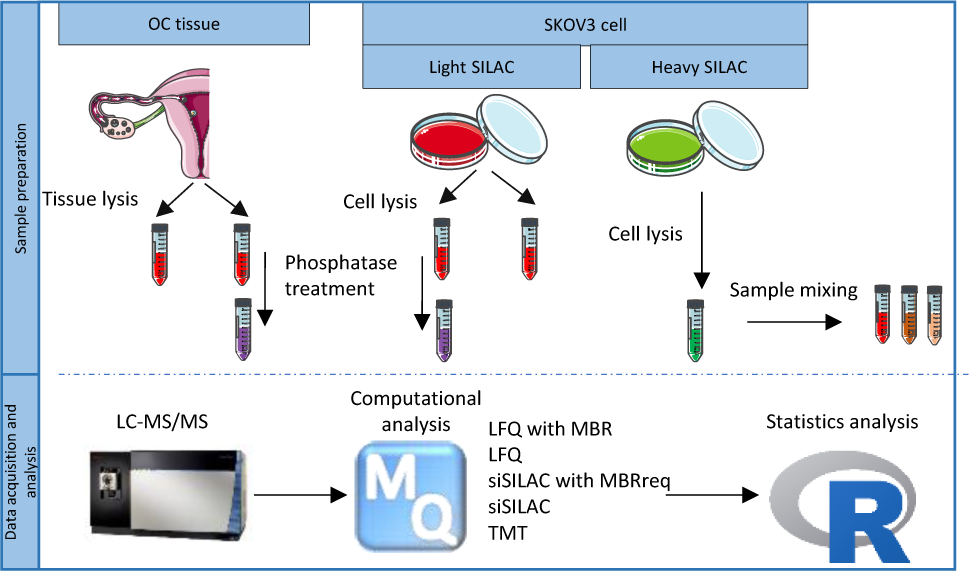

