## Supporting information (Supplementary figures, tables and experimental sections) for "Comparative assessment of quantification methods for tumor tissue phosphoproteomics"

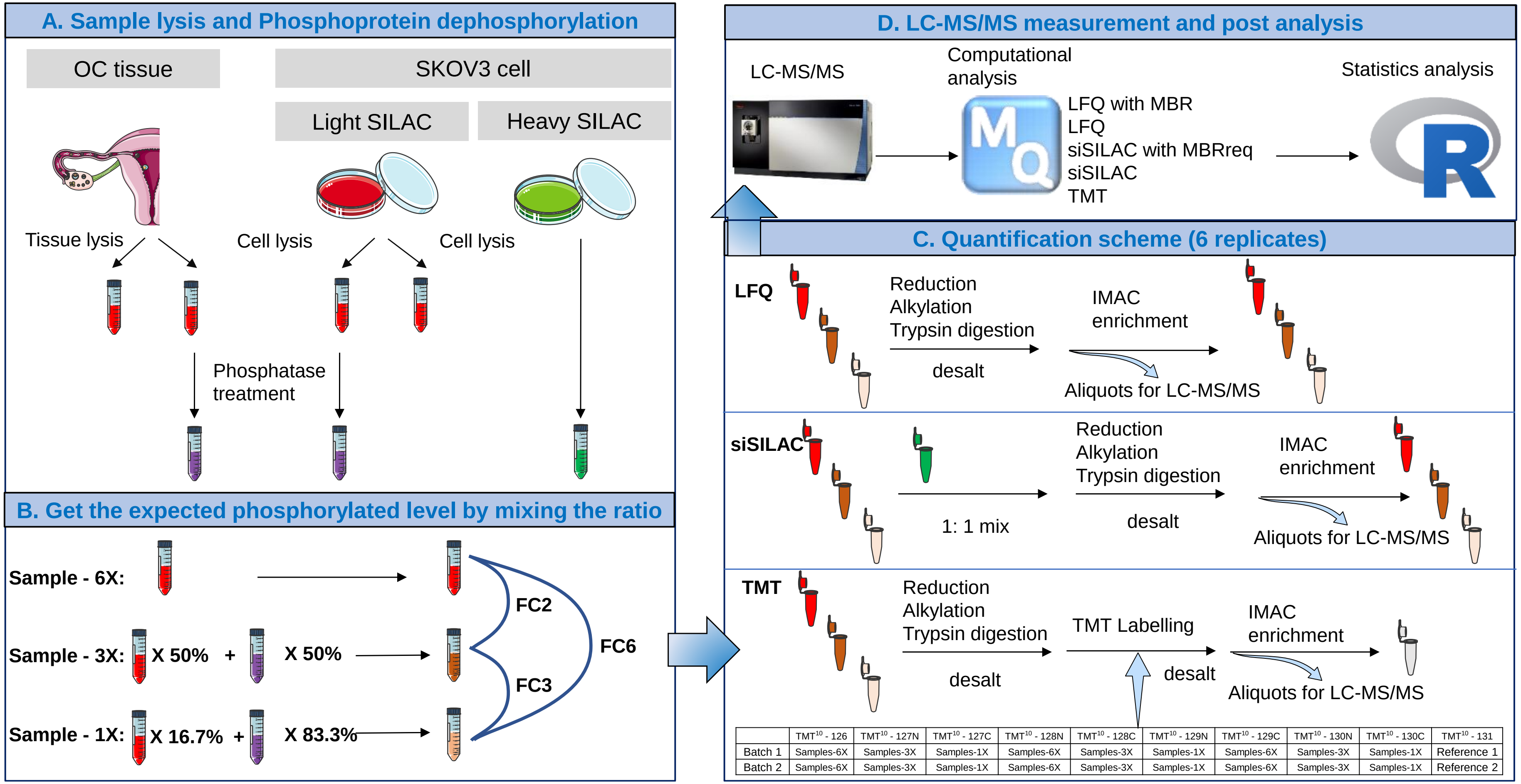


| | $TMT^{10}$ - 126 | $TMT^{10}$ - 127N | $TMT^{10}$ - 127C | $TMT^{10}$ - 128N | $TMT^{10}$ - 128C | $TMT^{10}$ - 129N | $TMT^{10}$ - 129C | $TMT^{10}$ - 130N | $TMT^{10}$ - 130C | $TMT^{10}$ - 131 |
|---|---|---|---|---|---|---|---|---|---|---|
| Batch 1 | Samples-6X | Samples-3X | Samples-1X | Samples-6X | Samples-3X | Samples-1X | Samples-6X | Samples-3X | Samples-1X | Reference 1 |
| Batch 2 | Samples-6X | Samples-3X | Samples-1X | Samples-6X | Samples-3X | Samples-1X | Samples-6X | Samples-3X | Samples-1X | Reference 2 |

**Figure S1: Study design: Sample preparation and MS measurement.** (Panel A) Proteins extracted from the tumor tissue and cell line were divided into two aliquots. One aliquot was dephosphorylated by alkaline phosphatase. To generate the spike-in-SILAC standard, the cells were cultured in heavy labelled SILAC media for 5 passages. (Panel B) To obtain samples with known phosphosite quantities for evaluating the quantitative performance, the original and dephosphorylated aliquots were mixed in a ratio of 1:5, 1:1 and 1:0, resulting in three sample groups with different phosphosite quantities (1X, 3X and 6X). The three sample groups allow the comparison of three different fold changes: 2, 3 and 6 (FC2, FC3, FC6). (Panel C) The samples were analyzed by LFQ, spike-in-SILAC and TMT 10 plex, with 6 technical replicates per sample. The TMT 10 plex included one reference sample for normalizing the batch variation. The reference consisted of a mixture of sample 6X, 3X and 1X in a ratio of 1:1:1. The scheme of sample labeling is shown in the table. (Panel D) The acquired raw data were processed with the MaxQuant suite, and the statistical analysis was done in R.

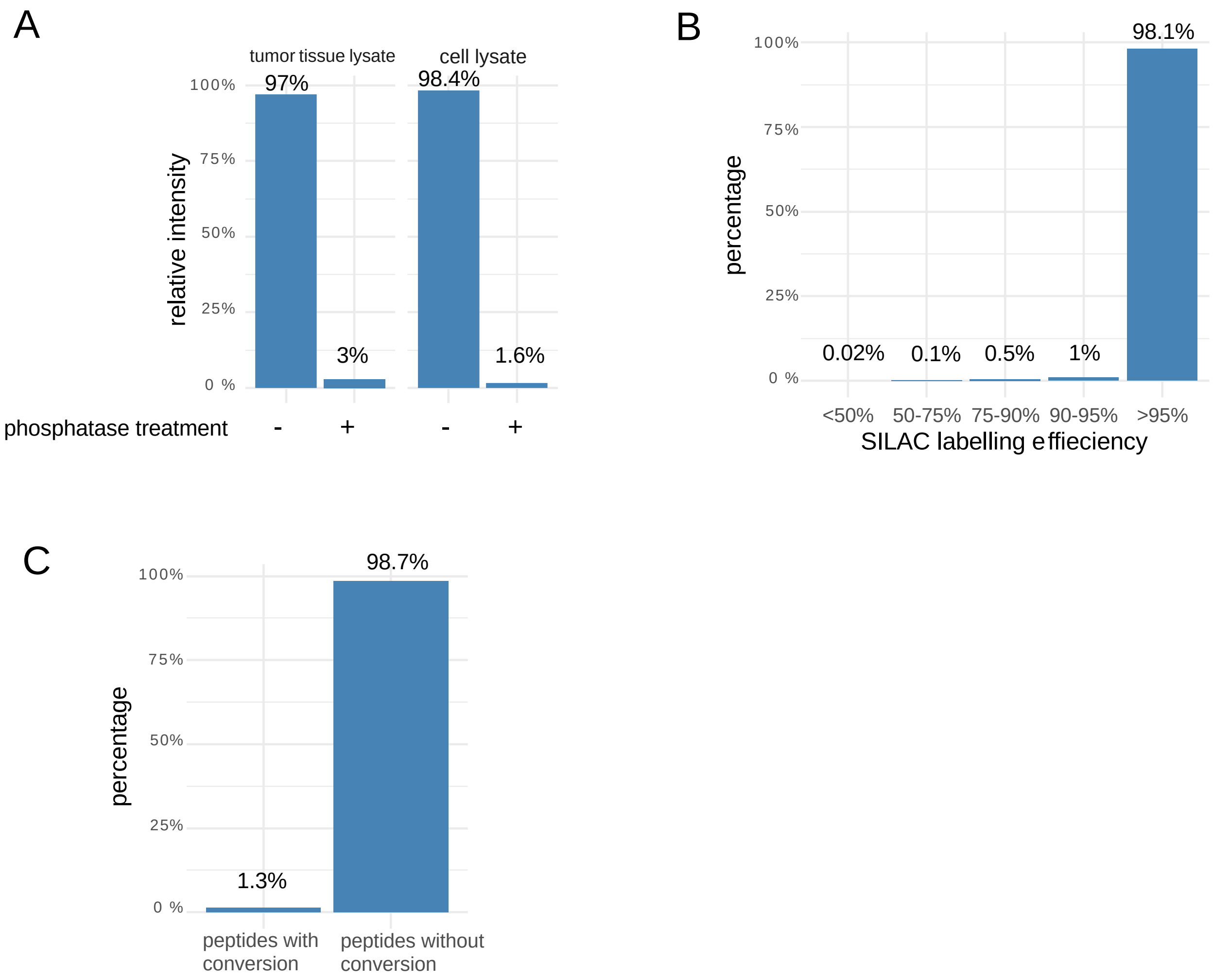


**Figure S2: Dephosphorylation efficiency, SILAC labelling efficiency and arginine-to-proline conversion.** (A) Relative intensities of identified phosphopeptides treated without (blue) and with (orange) alkaline phosphatase treatment in both tumor tissue lysate and cell lysate. (B) Bar plot showing the distribution of the identified peptides according to their SILAC labelling efficiency. (C) Relative intensities of peptides with and without arginine-to-proline conversion.

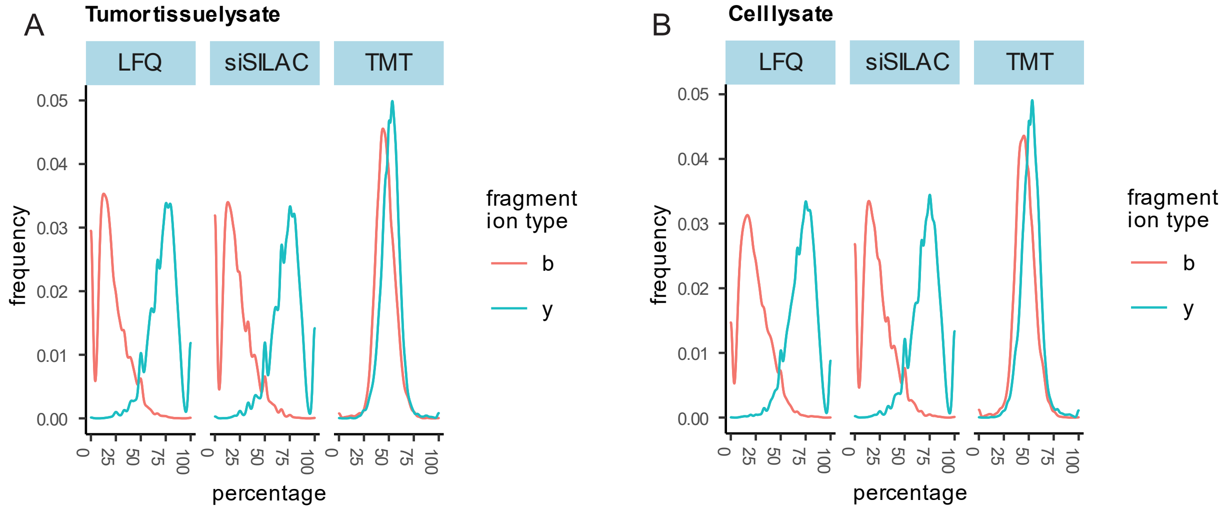


**Figure S3:** A, B: Density plots depicting the distribution of b- and y-ions of the identified peptides in tumor tissue lysates (A) and cell lysates (B).

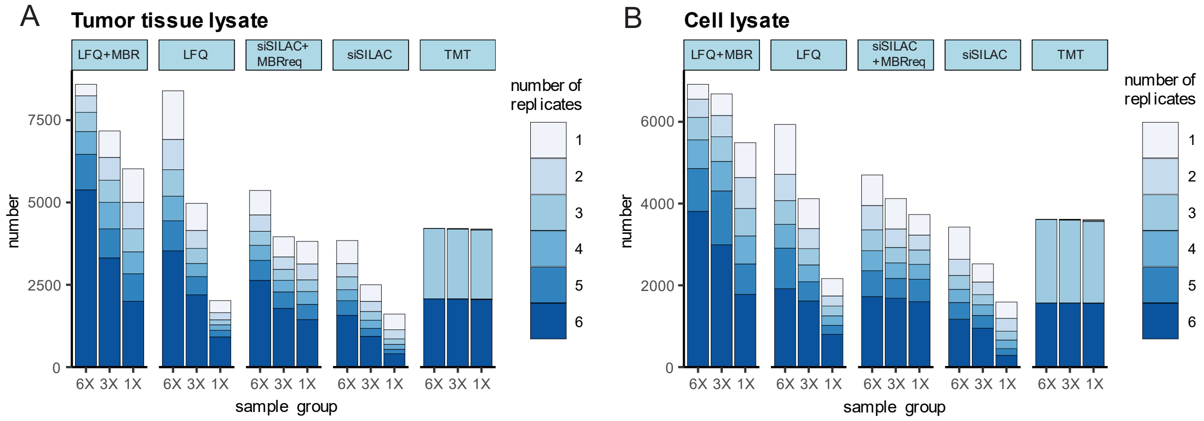


**Figure S4: Bar plots showing the number of phosphopeptides identified in each sample group (6X, 3X and 1X) in tumor tissue lysates (A) and cell lysates (B).** The color intensity indicates the number of replicates in which the phosphosites were identified.

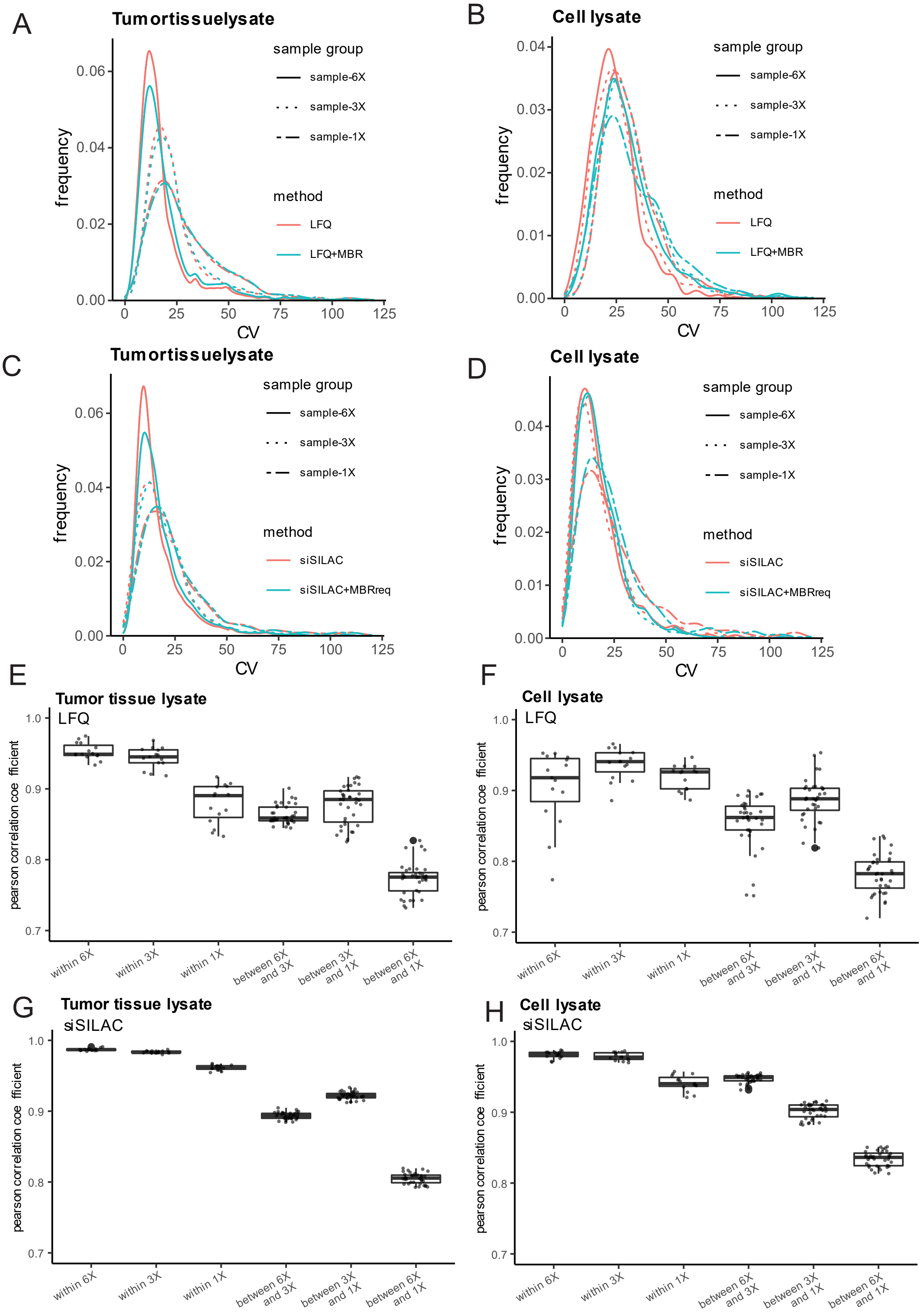


**Figure S5: Reproducibility of phosphosite quantification**. A, B: Density plots showing the distribution of CV values for phosphosites quantified with LFQ+MBR and LFQ in the tumor tissue lysates (A) and cell lysates (B). C, D: Density plots showing the distribution of CV values for phosphosites quantified with spike-in-SILAC + MBRreq and spike-in-SILAC in the tumor tissue lysates (C) and cell lysates (D). Different line types (solid, dotted, dashed) indicate different sample groups (6X, 3X and 1X). E-H: Boxplots visualizing the pairwise correlation coefficients within the same sample group (i.e. 6X vs. 6X, 3X vs. 3X, 1X vs. 1X) and between the different sample groups (i.e., 6X vs. 3X, 3X vs. 1X, 6X vs. 1X) in tumor tissue lysates (E: LFQ, G: spike-in-SILAC) and cell lysates (F: LFQ, H: spike-in-SILAC). The boxes show the first, second (median) and third quartile.

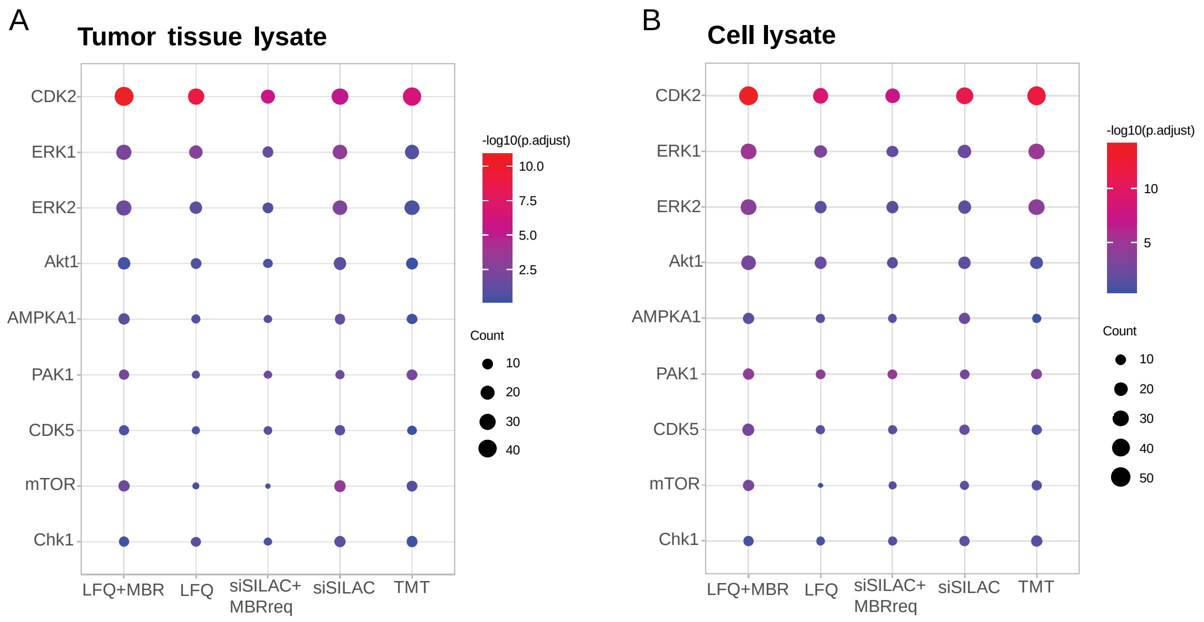


**Figure S6: Kinase substrate sites covered by the quantitative phosphoproteomes**. Dot-plots visualizing the enrichment analysis of quantified phosphosites from tumor tissue lysates (A) and cell lysates (B). The kinase-substrate database from PhosphositesPlus was used as background dataset. The size of the dots represents the number of phosphosites quantified for the corresponding kinase. The color intensity corresponds to the adjusted p-value using Benjamini-Hochberg approach.

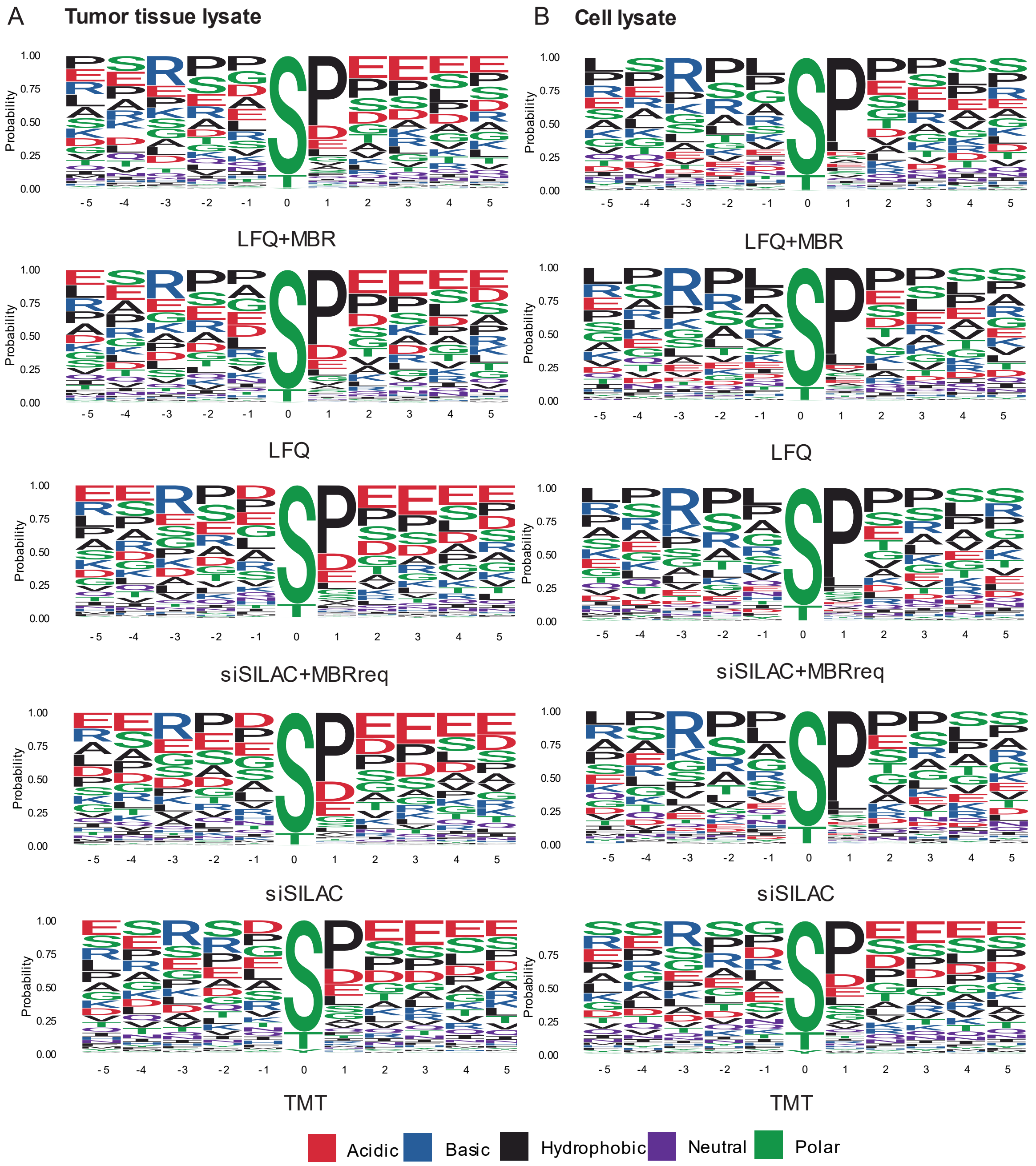


**Figure S7: Sequence motif analysis of quantified phosphosites from tumor tissue lysates (A) and cell lysates (B).** The phosphorylated residue is located at the central position within a 11-mer phosphorylated peptide sequence. The amino acids are colored according to their chemical properties and the size represents the observed probability.

### Supplementary tables

| | method | measurement time (hour) | number of MS2 | number of identified MS2 | number of MS3 | number of identified phosphopeptides | number of identified unique phosphopeptides (localization probability > 0.75) |
|---|---|---|---|---|---|---|---|
| tumor tisse lysate | LFQ | 36 | 471,216 | 171,753 | n.a | 8,932 | 5,812 |
| | siSILAC | 36 | 613,576 | 240,688 | n.a | 6,923 | 4,828 |
| | TMT | 4 | 63,421 | 8,122 | 63,345 | 4,281 | 3,578 |
| cell lysate | LFQ | 36 | 498,505 | 206,819 | n.a | 7,439 | 4,163 |
| | siSILAC | 36 | 597,456 | 236,793 | n.a | 5,881 | 3,621 |
| | TMT | 4 | 56,149 | 7,059 | 56,062 | 3,683 | 3,023 |

**Table S2: Key features of tumor tissue and cell phosphoproteomes quantified by LFQ, spike-in-SILAC (siSILAC) and TMT.** Shown are the measurement time, the number of MS2 spectra, the number of identified MS2 spectra, the number of MS3 spectra, the number of identified phosphopeptides and the number of identified phosphosites (minimum localization probability of 0.75).

| | **tumor tissue lysate** | | | | | |
|---|---|---|---|---|---|---|
| | true positive rate | | | AUROC | | |
| fold change | FC2 | FC3 | FC6 | FC2 | FC3 | FC6 |
| LFQ+MBR | 0.7500 | 0.9700 | 0.9900 | 0.9394 | 0.9852 | 0.9938 |
| LFQ | 0.9900 | 0.9940 | 0.9990 | 0.9953 | 0.9971 | 0.9990 |
| siSILAC+MBRreq | 0.7700 | 0.8800 | 0.9800 | 0.9442 | 0.9633 | 0.9899 |
| siSILAC | 0.9200 | 0.9700 | 0.9900 | 0.9702 | 0.9881 | 0.9960 |
| TMT | 0.9900 | 0.9970 | 0.9980 | 0.9942 | 0.9988 | 0.9990 |
| | **cell lysate** | | | | | |
| | true positive rate | | | AUROC | | |
| fold change | FC2 | FC3 | FC6 | FC2 | FC3 | FC6 |
| LFQ+MBR | 0.5990 | 0.9400 | 0.9690 | 0.8923 | 0.9749 | 0.9916 |
| LFQ | 0.5910 | 0.9200 | 0.9700 | 0.8992 | 0.9813 | 0.9900 |
| siSILAC+MBRreq | 0.9300 | 0.9000 | 0.9950 | 0.9817 | 0.9673 | 0.9931 |
| siSILAC | 0.9500 | 0.9800 | 0.9970 | 0.9815 | 0.9925 | 0.9988 |
| TMT | 0.9680 | 0.9930 | 0.9990 | 0.9879 | 0.9933 | 0.9985 |

**Table S3: True-positive-rates (TPRs) for the different quantification methods** at a false-positive-rate (FPR) threshold of 0.05, and the area under the receiver operating characteristic (AUROC).

### Supplementary experimental section

**SKOV3 cell culture and SILAC labeling.** The SKOV3 cells were purchased from ATCC (HTB-77, Homo sapiens ovary). Cell were cultivated in lysine- and arginine-free Dulbecco's modified Eagle's medium (1111DMEM) with 1 g/L glucose, (PAN Biotech, cat. no. P04-02506), supplemented with 10% (v/v) dialyzed fetal bovine serum (FBS) (Gibco, cat. no. 10270-106), D-(+)-Glucose solution (Sigma-Aldrich, cat. no. G8644-100 ML) to a final concentration of 4.5 g/L, 3 mM L-glutamine (Gibco, cat. no. 25030-024), 84 mg/L lysine and 146 mg/L arginine. For SILAC labeling "heavy" lysine ($^{13}C6$, $^{15}N2$, Cambridge Isotope Laboratories Inc., CNLM-291-H-PK) and arginine ($^{13}C6$, $^{15}N4$, Cambridge Isotope Laboratories Inc., CNLM-539-H-PK) (heavy labeled medium) or "light" lysine (Cambridge Isotope Laboratories Inc., ULM-8766-PK) and arginine (Cambridge Isotope Laboratories Inc., ULM-8347-PK)[1] (light labeled medium) was used. For TMT and LFQ analysis cells cultivated in light labeled medium were used. All cells were maintained at 37°C and 7.5% $CO_2$ for at least five passages and regularly tested for mycoplasm. Arginine-to-proline conversion and SILAC labeling efficiency was tested by LC-MS/MS. The arginine-to-proline conversion was at or below 1% and the SILAC labeling efficiency was higher than 98%.

**SKOV3 cell line and ovarian cancer tissue lysis**. For serum and amino acid starvation, SKOV3 cells were washed with phosphate-buffered saline (PBS, PAN Biotech, cat. no. P0436500) and cultured for 16 h in amino acids free DMEM medium (PAN Biotech, cat. no. P0401507). For stimulation, the medium was exchanged to light labeled medium for 90 min. Cells were washed three times with ice-cold PBS and lysed in protein lysis buffer [8 M urea, 75 mM NaCl, 50 mM Tris, pH 8.2] supplemented with Complete Mini (Roche, cat. no. 04 693 124 001) and PhosSTOP (Roche, cat. no. 04 906 845 001).

The ovarian cancer tissue was provided by PROMETOV and analyzed under the ethics board approval S496/2014. For tissue lysis, frozen tumor samples were crushed on dry ice and subsequently pulverized at 35 Hz for 2 min in liquid nitrogen precooled teflon milling cups using a ball mill (Retsch Mixer Mill MM 400) and a metal ball. The well-mixed tissue powder was aliquoted on dry ice before taking up in protein lysis buffer.

Cell lysates and tissue lysates were sonicated at 20% amplitude with a 20 x 1 sec pulse on ice (1 sec sonication and 1 sec cooling down), before centrifugation at 12,500 × g, 4°C for 10 min. The supernatant was transferred to new tubes. The protein concentrations were measured using the BCA Protein Assay Kit (Thermo-Fisher Scientific, cat.no. 23235) according to the manufacturer's protocol and adjusted to 1 mg/mL.

**Protein reduction, alkylation and digestion.** To reduce disulfide bonds, lysates were incubated with 5 mM DTT at 37°C under gentle shaking for 1 h. To alkylate cysteine residues, iodoacetamide was added to a final concentration of 14 mM and incubated for 30 min at RT in the dark. Subsequently, DTT was added to a final concentration of 5 mM and the samples were incubated for an additional 15 min at RT in the dark. Next, the samples were diluted with a ratio of 1:5 with 25 mM Tris-HCl, pH 8.2 to reduce the urea concentration to 1.6 M. For tryptic digestion, the samples were incubated with trypsin (trypsin to substrate ratio of 1:50, Promega, cat. no. V5111) and 1 mM $CaCl_2$ for 16 h at 37°C. The tryptic digestion was stopped by adding TFA to a final concentration of 0.4% (v/v), respectively. The samples were desalted using SepPak tC18 cartridges (Sep-Pak tC18 1 cc Vac Cartridge, 100 mg Sorbent per Cartridge, 37 - 55 µm, cat. no. WAT036820). The cartridges were conditioned using two times 1 mL condition buffer (95% ACN, 5% $H_2O$, 0.1% TFA), before washing two times with 1 ml of wash buffer (5% ACN,95% $H_2O$, 0.1% TFA). Samples were loaded to the cartridges followed by washing three times with 1 mL wash buffer. The peptides were eluted using two times 1 mL elution buffer (50% ACN, 50% $H_2O$). For the LFQ and spike-in-SILAC measurements, 50 µL of the eluent was used for the LC-MS/MS analysis and 950 µL for the subsequent IMAC enrichment. All the samples were dried using a SpeedVac.

**LC-MS/MS measurement.** For LC-MS/MS analysis, samples were injected on an ultrahigh performance nano liquid chromatography system (Dionex UltiMate 3000 RSLCnano, Thermo Scientific, Bremen, Germany) coupled to an Orbitrap mass spectrometer (Fusion™, Thermo Fisher Scientific) with a nano electrospray source.

The samples were loaded (3 µL/min) with the buffer A (0.1% formic acid (FA) in HPLC grade $H_2O$) on a trapping column (Acclaim PepMap µ-precolumn, C18, 300 µm × 5 mm, 5 µm, 100 Å, Thermo Scientific, Bremen, Germany). After sample loading, the trapping column was washed with 30 µL buffer A (3 µL/min) and the peptides were eluted (300 µL/min) onto separation column (Acclaim PepMap 100, C18, 75 µm × 500 mm, 2 µm, 100 Å, Thermo Scientific, Bremen, Germany). The column temperature was kept constant at 45°C and the peptides were separated with a gradient from 5−25% buffer B in 90 min. The spray was generated from a silica emitter with conductive coating (O.D. 360 µm, I.D. 20 µm, Tip I.D. 10 µm New Objective, Littleton, USA) at a capillary voltage of 1800 V. MS analyses were performed in positive ion mode and data-dependent acquisition mode (DDA). LC-MS/MS analysis was carried out in a cycle time of 3 s and the dynamic exclusion duration was set to 30 s. For the spike-in-SILAC and LFQ samples, an intensity threshold of $2{\cdot}10^4$, an isolation width of 0.7 m/z and an HCD collision energy of 30 NCE was used for MS/MS experiments. Precursor MS scans were performed over a m/z range from 380-1500, with a resolution of 120,000 FWHM at m/z 200 (RF Lens = 60%, maximum injection time= 50 ms, AGC target= $2{\cdot}10^5$) and MS/MS spectra were recorded with a resolution of 7,500 FWHM at m/z 200 (maximum injection time= 22 ms, AGC target= $2{\cdot}10^5$).

For TMT, we used multi-notch synchronous precursor selection (SPS)-MS3 technology to co-isolate and co-fragment multiple MS2 fragment ions ion the linear ion trap, followed by reporter ions detection in the orbitrap (MS3). The SPS-MS3 approach reduces reporter ion variance and enhance quantification accuracy[2,3]. An intensity threshold of $5{\cdot}10^3$ and an isolation width of 0.7 m/z was used for MS/MS experiments. Precursor MS scans were performed over a m/z range from 380-1500, with a resolution of 120,000 FWHM at m/z 200 (RF Lens = 60%, maximum injection time= 50 ms, AGC target = $2{\cdot}10^5$) and the precursors were fragmented by collision-induced dissociation (CID) (AGC $1{\cdot}10^5$, normalized collision energy (NCE) = 35, q-value = 0.25, maximum injection time = 50 ms, isolation window = 0.7) and measured in the ion trap (ion trap scan rate = turbo). For each MS2 spectrum, we collected an MS3 spectrum in which multiple MS2 fragment ions are captured using isolation waveforms with multiple frequency notches. MS3 precursors were fragmented by HCD and analyzed using the Orbitrap (NCE = 65, AGC = $1{\cdot}10^5$, maximum injection time = 120 ms, resolution = 60,000, m/z range = 120-500).

**Raw data processing.** All LC-MS/MS data were processed with MaxQuant[4] version 1.6.5. Peptides and proteins identification was performed using the Andromeda search engine[5] and searched against the Swissprot database (Uniprot, downloaded 2019-09-20, 20430 entries) and a contamination database (cRAP-database, http://www.thegpm.org/crap, 298 entries). For analysis of LFQ measurements, data were searched with and without “match between runs” (MBR). For analysis of the spike-in-SILAC measurements, the data were searched with and without MBR and re-quantify (req) option. TMT correction factors were added in MaxQuant using the values provided by the manufacturer. Carbamidomethylation of cysteines was specified as fixed modification and oxidation of methionine, and N-terminal protein acetylation and phosphorylation of serine, threonine and tyrosine residues were defined as variable modifications. Only phosphopeptides with a localization probability larger than 0.75 were considered for further analysis.
